# 4-Aminopyridine Produces Concentration-Dependent Improvement in Locomotor Recovery after Spinal Cord Injury in Zebrafish Larvae^1^

**DOI:** 10.1101/2024.07.15.603582

**Authors:** Payge M. Hoffman, Natalie L. Clark, Karen Mruk

## Abstract

Spinal cord injury (SCI) affects 250,000 to 500,000 individuals annually. After the initial injury, a delayed secondary cascade of cellular responses occurs causing progressive degeneration and permanent disability. One part of this secondary process is disturbance of ionic homeostasis. The K^+^ channel blocker, 4-aminopyridine (4-AP), is used clinically to alleviate symptoms of multiple sclerosis (MS). Several ongoing studies are being conducted to explore additional areas where 4-AP may have an effect, including stroke, traumatic brain injury, and nervous system recovery after SCI. The goal of our study was to determine whether 4-AP affects functional recovery from SCI in zebrafish (*Danio rerio*). Using the transgenic line *Tg(gfap:EGFP; elavl3:mCherry-CAAX)*, we created a spinal transection, tracked swim recovery, and measured 4-AP toxicity. We found that constant treatment with 10 µM 4-AP increases swimming distance 40%. Live imaging demonstrated that treatment with 4-AP did not accelerate axonal bridging but did affect radial glial cells suggesting that 4-AP affects functional recovery through mechanisms other than axon regeneration. Higher concentrations provided no additional functional benefit and led to increased toxicity in zebrafish.

**Significance Statement:** In this study, we found that 4-AP can enhance locomotor recovery in zebrafish after spinal cord injury. Our findings indicate that inhibition of K^+^ channels with 4-AP may promote glial remodeling that is pro-regenerative.

## Introduction

Approximately 17,000 people in the United States suffer a spinal cord injury (SCI) each year (2017). Many complications arise with SCI, such as cardiovascular and pulmonary effects, along with the expected neurological effects. Immediate interventions, such as decompressive surgery, are used to alleviate and/or avoid secondary complications like hypoxia and ischemia. However, management of chronic SCI involves treating the secondary complications, with very few pharmacological treatments for promoting regeneration.

4-aminopyridine (4-AP) is an agent that antagonizes voltage-gated potassium (K_V_) channels (Gillespie 1977) and is therapeutically used in various demyelinating disorders (Hayes 2004). Nonspecific antagonism of K_V_ channels enhances axonal conduction and prolongs the action potential of damaged, demyelinated axons. 4-AP is currently available for prescription as symptomatic relief for individuals with multiple sclerosis (Dietrich et al. 2021; Goodman and Stone 2013; Leussink et al. 2018). Increased impulse conduction caused by K_V_ channel inhibition has been shown to improve walking speed and lower extremity muscle strength (Jensen et al. 2014). More recently, studies have demonstrated that 4-AP prevents axonal loss in addition to relieving the symptoms of MS (Dietrich et al. 2020). Furthermore, K_V_ channels are expressed in innate immune cells and modulatation of these channels inhibits immune cell proliferation and proinflammatory cytokine release (Dietrich et al. 2021; Villalonga et al. 2010; Wang et al. 2019; Wulff et al. 2003).

In SCI, multiple cell types are affected including: neurons, oligodendrocytes, glial cells, and innate immune cells. Systematic review of randomized clinical trials indicate that 4-AP may improve function in patients with SCI, depending on whether the spinal tract is preserved and the extent of myelination (Paredes-Cruz et al. 2022; Wiener et al. 2018). Therefore, we sought to determine whether 4-AP could enhance functional recovery after a complete spinal cord transection. We used the pro-regenerative animal model zebrafish (*Danio rerio)* because zebrafish have a well-defined locomotor repertoire. Zebrafish locomotion recovers after a complete spinal transection, making this model ideal to answer questions about the effectiveness of 4-AP without interference from spared tissue. 4-AP has previously been used in zebrafish larvae to induce hyperactivity at concentrations above 500 micromolar (Ellis et al. 2012). We now show that low concentrations of 4-AP can enhance locomotor recovery after a spinal cord transection in zebrafish larvae.

## Materials and Methods

### Zebrafish husbandry

Adult transgenic zebrafish *Tg(gfap:EGFP)* 3-18 months were housed in our laboratory. Adult *Tg(gfap:EGFP; elavl3:mCherry-CAAX)* were generated by our laboratory as reported previously (Walker et al. 2025). Adults were maintained at 28.5°C in a recirculating system (Iwaki Aquatic) on a 14:10 hr light:dark cycle. Adults were fed once with Zeilger’s adult zebrafish diet (Pentair Aquatic Ecosystems) and once with brine shrimp (E-Z Egg, Brine Shrimp Direct). Embryos were obtained through natural matings. Embryos were cultured at 28–30°C in E3 medium containing 5 mM NaCl, 0.17 mM KCl, 0.33 mM CaCl2, and 0.33 mM MgSO4. All chemicals were purchased from Sigma Aldrich. Embryos were staged as described previously (Kimmel et al. 1995). Zebrafish larvae began receiving rotifers (Reed Mariculture) at five days post fertilization (dpf) unless otherwise noted. The IACUC committees at the University of Wyoming and East Carolina University approved all animal procedures.

### Spinal Cord Transection

5 dpf larvae were mounted on flat slides in 2% low melting point (LMP) agarose and anesthetized with 15 μM of 0.01% Syncaine/Tricaine-S (MS-222, Syndel). After anesthetization, each larvae was transected at the level of the cloaca using a 5 mm microsurgical blade (World Precision Instruments). Following transection, larvae were imaged to confirm a complete transection (Supplementary Fig. 1). Larvae recovered in individual wells of a 48-well plate containing 1 mL of 1x E2 working solution supplemented with penicillin/streptomycin (Pen/Strep 100 unit, 100 µg/mL, Gibco #15140-122) and kept at 28°C. E2 media was replaced daily with fresh antibiotics. Larvae were not fed on day of injury to reduce risk of infection.

### Treatment with 4-AP

4-AP was purchased from Sigma and made into a stock concentration of 100 mM dissolved in dimethyl sulfoxide (DMSO). After spinal transection at 5 dpf, larvae were placed in individual wells on a 48-well plate, each well containing 1x E2 working solution with penicillin/streptomycin and either 10, 20, 40, or 100 μM of 4-AP or an equal volume of DMSO. An equal number of larvae was used for each of the following conditions: uninjured DMSO, uninjured 4-AP, spinal transection DMSO, and spinal transection 4-AP. The plate was kept at 28° C overnight. Each day, prior to behavioral recording, a fresh plate was prepared with new E2 working solution with penicillin/streptomycin and either 4-AP or DMSO and the larvae were transferred to the new plate prior to being placed on the Zebrabox for one hour. Behavior was not tracked on day of injury. After completion of each behavioral recording, larvae were fed with a diluted rotifer culture such that each well had ∼100 rotifers / mL.

### 4-AP toxicity

Control and injured larvae were imaged daily on an Olympus SZX16 stereomicroscope equipped with an Olympus DP80 camera and SDF PLANO 1XPF objective or a Leica M165FC and 1x M-series Planapo objective equipped with a Leica K7 camera. The pre-installed CellSens software (Olympus Life Science) or LasX software (Leica) was used to capture and save images. Survival was recorded on a daily basis. Images were opened in FIJI open-source software (Schindelin et al. 2012). The following parameters were measured to assess toxicity: total length, swim bladder area, eye area, and liver intensity.

### Imaging and fluorescence intensity quantification

Larvae were transected as described above. Both transected and uninjured controls were mounted in 1% LMP agarose and imaged on a Leica DM6D epifluorescence microscope with a Leica K8 camera and CoolLED pE-300 Ultra light source 20x/NA 0.5 water immersion objective. Z-stacks were generated from images taken at 2-5 µm intervals. To quantify pixel intensity, fluorescent 20x z-stacks were opened in FIJI as maximum projections. The region of interest was selected using the segmented line selection tool at 200-point width. The Plot Profile analysis tool was then used to measure pixel intensity across the region of interest. Larvae with injuries larger than 250 microns and larvae that did not survive the entire study were excluded from analysis.

### Swimming behavior

Larvae were transected as outlined above and their behavior was recorded for seven days and compared to normally developing larvae. Larval zebrafish reliably show activity on transition to a dark environment from a lit environment and are more active in the morning beginning around 10 am and reach their lowest activity in the evening by 5 pm (MacPhail et al. 2009). Behavior experiments were executed between 14:00 and 17:00 local time to ensure reproducibility. A clear 48-well plate containing 1 mL of fresh E2 supplemented with Pen/Strep was used for experiments, with one larva per well. Locomotion was assessed using ZebraBox and its ViewPoint LS Tracking Software (v5.15.0.230, ViewPoint Life Sciences, Lyon, France). Larvae were acclimated to the test environment for 10 minutes, followed by a 10-minute session in a dark environment and behavior was video recorded and tracked every 10 minutes for 60 minutes. Total swim distance was found by adding the tracked distances together. The movement speed thresholds were set to: 0-2 cm/s, 2-4 cm/sec, >4 cm/sec. Following the last day of behavior experiments, larvae were imaged as outlined above to record the absence or presence of a glial bridge.

### Statistical Analysis

For all experiments, breeding tanks with 2 to 3 males and 3 to 4 females from the *Tg(gfap:EGFP)* or *Tg(gfap:EGFP; elavl3:mCherry-CAAX)* lines were set up to generate embryos. Collected embryos were randomly distributed for all testing conditions. Developmentally abnormal or unfertilized embryos were removed before transection or initiation of treatment. Prior to all experimentation, it was decided that larvae that did not survive the entire course of the experiment were to be excluded from all behavioral and imaging analyses. Sample sizes were calculated to observe a 30% effect with 80% power. Graphs were plotted using GraphPad Prism. All swim data was analyzed by a Shapiro-Wilk test for normality and determined to be non-normal. Significant differences in swim behavior were calculated using a mixed-effects analysis using the Geisser-Greenhouse correction with post-hoc Tukey multiple comparison test. Significant differences in survival curves was determined using log-rank test Mantel-Cox.

## Results

### 100 μM 4-AP causes toxicity in 5 dpf sci zebrafish larvae

Previous research suggested that 4-AP concentrations of 2.5 mM 4-AP eliminated fast movement in acute treatment (Ellis et al. 2012). Our aim was to find therapeutic concentrations that would help recover and maintain fast, deliberate movement. We determined a beginning dose of 100 µM according to the recommended human dose of 0.5 mg/kg (Van Diemen et al. 1993). After 5 dpf, larvae were completely transected as verified by EGFP imaging post-injury. This line labels radial glial throughout the spinal cord of the zebrafish, acting as a good marker to report on both transection and formation of a pro-regenerative glial bridge after SCI. All transected larvae incubated with 100 µM 4-AP had a significant decline to 0% survival at 4 days post injury (dpi) (Fig. [1A]) compared to those treated with constant 0.1% DMSO (P value < 0.0001). After two repeated trials with 0% survivability of transected 100 µM 4-AP treated embryos, one trial of uncut control fish was treated with 100 µM 4-AP to determine whether this concentration was toxic to zebrafish larvae independent of a spinal transection. The uncut control larvae also experienced a significant decline to 0% survival at 4 dpi (P value < 0.0001). We next aimed for shorter exposure times. Daily 3 hr treatment of 100 µM 4-AP did prolong lifespan but ultimately led to complete mortality with no significant improvement of locomotor recovery (Supplemental Fig. 2).

Because 100 µM 4-AP treated larvae had a 0% survival rate, we decreased the dose 10-fold to 10 µM for constant treatment over the course of seven days. After transecting 5 dpf *Tg(gfap:EGFP)* larvae, fish were incubated in either constant 10 µM 4-AP or an equal volume of DMSO, the vehicle control. Larvae treated with 10 µM 4-AP had a 90% survival rate at 4 dpi. At 7 dpi, the survival rate of 10 µM treated larvae was 80.6% and 0.1% DMSO treated larvae was 80.3% (Fig. [1B]). In addition, the survival of uncut 10 µM 4-AP treated larvae was 97% at 7 dpi, indicating constant exposure to this dose of 4-AP was not toxic to zebrafish larvae.

### 4-AP increases locomotor recovery in transected zebrafish larvae greater than the vehicle control counterpart

A previous study of intrathecal delivery of 4-AP to patients with chronic spinal cord injury found some recovery of sensory function and spasticity in a small percentage of patients in the study (Halter et al. 2000). Similarly, recovery of sensory and motor function was also reported in patients taking oral 4-AP for 12 weeks compared to placebo (Grijalva et al. 2003). Therefore, we next determined whether 4-AP would increase motor function in zebrafish larvae. After confirmed transection of 5 dpf *Tg(gfap:EGFP)* larvae, fish were treated with either 10 µM 4-AP or 10 µM DMSO. Individual fish were tracked over the course of seven days (Fig. [2A]). Beginning at 3 dpi, there was a significant difference in total distance moved between 4-AP treated and DMSO treated SCI fish. 4-AP treated larvae continued to outswim DMSO-treated larvae throughout the rest of the study.

Unlike mammals which form an inhibitory glial scar, zebrafish form a pro-regenerative glial bridge after SCI (Briona and Dorsky 2014; Goldshmit et al. 2012; Klatt Shaw et al. 2021; Mokalled et al. 2016; Purifoy and Mruk 2024). This bridge is required to permit axons to cross the injury site (Zhou et al. 2023). Imaging of *Tg(gfap:EGFP)* larvae at the beginning and end of the behavioral experiment, indicated differences in glial bridging at the site of injury (Fig. [2B]). Therefore, we next used our *Tg(elavl3-mCherry-CAAX; gfap:EGFP)* larvae to quantify glial intensity and glial and axonal bridging. Although the total percentage of zebrafish that formed a glial bridge was only slightly increased in 4-AP treated animals, (92% vs 88%), there was less accumulation of GFP signal around the site of injury in 4-AP treated larvae (Fig. [3A,B]). Treatment with 10 µM 4-AP did not increase the amount of axonal bridging (Fig. [3C]). Fitting the data to a logistic growth model indicated that larvae treated with 4-AP formed an axonal bridge with slightly slower kinetics (τ_DMSO_ – 12.9 h, τ_4-AP_ – 15.4 h).

### Increased 4-AP concentrations do not enhance swim recovery

One of our objectives was to determine an ideal dose to optimize swim recovery while obtaining a high survival rate. We choose two increased concentrations, 20 and 40 µM. After transection of 5 dpf larvae, fish were continuously incubated with 20 µM 4-AP, 40 µM 4-AP or an equal volume of DMSO. Compared to the transected DMSO controls, there was no significant difference in survival rate of transected 20 µM 4-AP treated larvae with rates of 83.3% and 77% respectively (Fig. [4A]). In addition, no noticeable differences were observed in the injury site; however, increased density in the liver was observed in surviving larvae (Supplemental Fig. 3). We measured swim distance in uncut and transected larvae for a full week after injury. No significant differences between 20 µM 4-AP and DMSO treated larvae were observed. Transected larvae treated with 40 µM 4-AP did have a significant reduction in survival at 7 dpi. (p value = 0.0006) (Fig. [4B]). Similar to 20 µM 4-AP treated larvae, surviving fish exposed to higher concentrations had no obvious changes in the site of injury, an increase in liver density, and no improvement in swimming.

## Discussion

We show that continuous exposure to 10 µM 4-AP increases the rate of swim recovery after SCI. Our findings in zebrafish larvae present a new model for testing the toxicity and efficacy of compounds and complement extensive work in mammals regarding 4-AP and improved locomotor function. The fact that we found increased recovery of movement in an already regenerative model highlights the utility of zebrafish larvae as a model to screen potential therapeutic compounds. Previous studies using 4-AP in zebrafish have focused on its use as a method of chemical induction of seizure (Williams and Mruk 2022). Zebrafish tolerate high transient doses (0.6 – 2.5 mM) of 4-AP in zebrafish but these doses induce anxiety/stress type behaviors to slow circling behavior in a concentration-dependent manner (Ellis et al. 2012). Even in our highest continuous dose, 100 µM, we did not observe these complex swimming behaviors.

Despite the lack of stress type swimming, higher doses did result in increased mortality coupled with increased density in the liver. Although 1% DMSO is typically cited as the maximal tolerated concentration, we also saw increased liver density in our 0.1% DMSO-treated larvae. There is limited metabolism of 4-AP in humans with ∼90% excreted unchanged in the urine. However, 4-AP has been shown to inhibit CYP2E1 with an IC50 value of 125 μM (Caggiano and Blight 2013). Zebrafish have two homologous P450 enzymes, Cyp2y3 and Cyp2p6 (Tsedensodnom et al. 2013). Given that outgrowth of the liver is considered complete in zebrafish at 5 dpf (Chu and Sadler 2009), our results suggest that larval zebrafish may afford a new platform to screen for potential cytochrome P450 inhibition and/or liver toxicity. Additional studies investigating the expression levels of specific Cyp enzymes are warranted.

Given the increase in swim behavior was not reminiscent of hyperactivity models, we hypothesized that the observed swim recovery was based on regeneration of normal circuitry. Our data suggest that treatment with 10 μM 4-AP does not substantially affect the timing or extent of axonal bridging following spinal cord injury. 4-AP may be improving conduction through injured or actively regenerating axons to promote recovery of swimming behavior.

We also looked at the radial glia population of cells in the spinal cord. Our lab and others have reported the formation of a glial bridge after SCI in larval zebrafish larvae beginning 2–3 dpi (Briona and Dorsky 2014; Purifoy and Mruk 2024; Wehner et al. 2017). In adults, regenerating axons associate with the elongated glia to cross the site of injury (Goldshmit et al. 2012; Mokalled et al. 2016). Larval zebrafish also form a glial bridge though the requirement of bridge formation for axon regrowth is still debated (Wehner et al. 2017; Zhou et al. 2023). Unexpectedly, animals treated with 4-AP had reduced GFP signal compared to DMSO-treated controls. 4-AP could be working directly on astroglia, either affecting proliferation at the site of injury or migration to the site of injury. Alternatively, changes in the excitability of surviving neurons due to 4-AP block may be creating a localized ionic gradient that affects astroglial movement.

Spinal cord astrocytes in mammals exhibit four distinct K^+^ currents: K_IR_, K_DR_, K_A_, and K_Sl_ that are selectively inhibited by 4-AP (Bordey and Sontheimer 1999). K^+^ channels in astrocytes play a critical role in clearing K^+^ by absorbing excess extracellular K^+^ and redistributing it to sites of low extracellular K+ (aka spatial buffering) (Murakami et al. 2015). In mammals, SCI produces a large and rapid rise in K+ at the injury site. Additional studies focused on spinal radial glial cells in zebrafish larvae provide an opportunity to better understand differences in ionic changes after SCI and how they contribute to functional recovery.

## Acknowledgements

We gratefully acknowledge financial support from NIH P20GM121310 and R03NS136719 (K.M.). We thank Thomas Rynes for excellent fish care. We thank members of the Mruk lab for helpful discussions and edits to the manuscript.

## Data Availability

The authors declare that the data supporting the findings of this study are available within the paper and its Supplemental Data. Any data not included are available upon request from the corresponding author.

## Author Contributions

*Participated in research design:* Hoffman, PM and Mruk, K

*Conducted experiments:* Hoffman, PM, Clark, NL, and Mruk, K

*Performed data analysis:* Hoffman, PM, Clark, NL, and Mruk, K

*Wrote or contributed to the writing of the manuscript:* Hoffman, PM, Clark, NL, Mruk, K

## Footnotes

This work was supported by the National Institutes of Health (NIH) National Institute of Neurological Disorders and Stroke [Grant R03NS136719] and National Institutes of Health (NIH) National Institute of General Medical Sciences [Grant P20GM121310].

## Abbreviations

(4-AP): 4-aminopyridine
(dpf): days post fertilization
(dpi): days post injury
(DMSO): dimethyl sulfoxide
(EGFP): enhanced green fluorescence protein
(MS): multiple sclerosis
(SCI): spinal cord injury
(K_V_) channel: voltage-gated potassium

## Figure Legends

**Figure 1.**
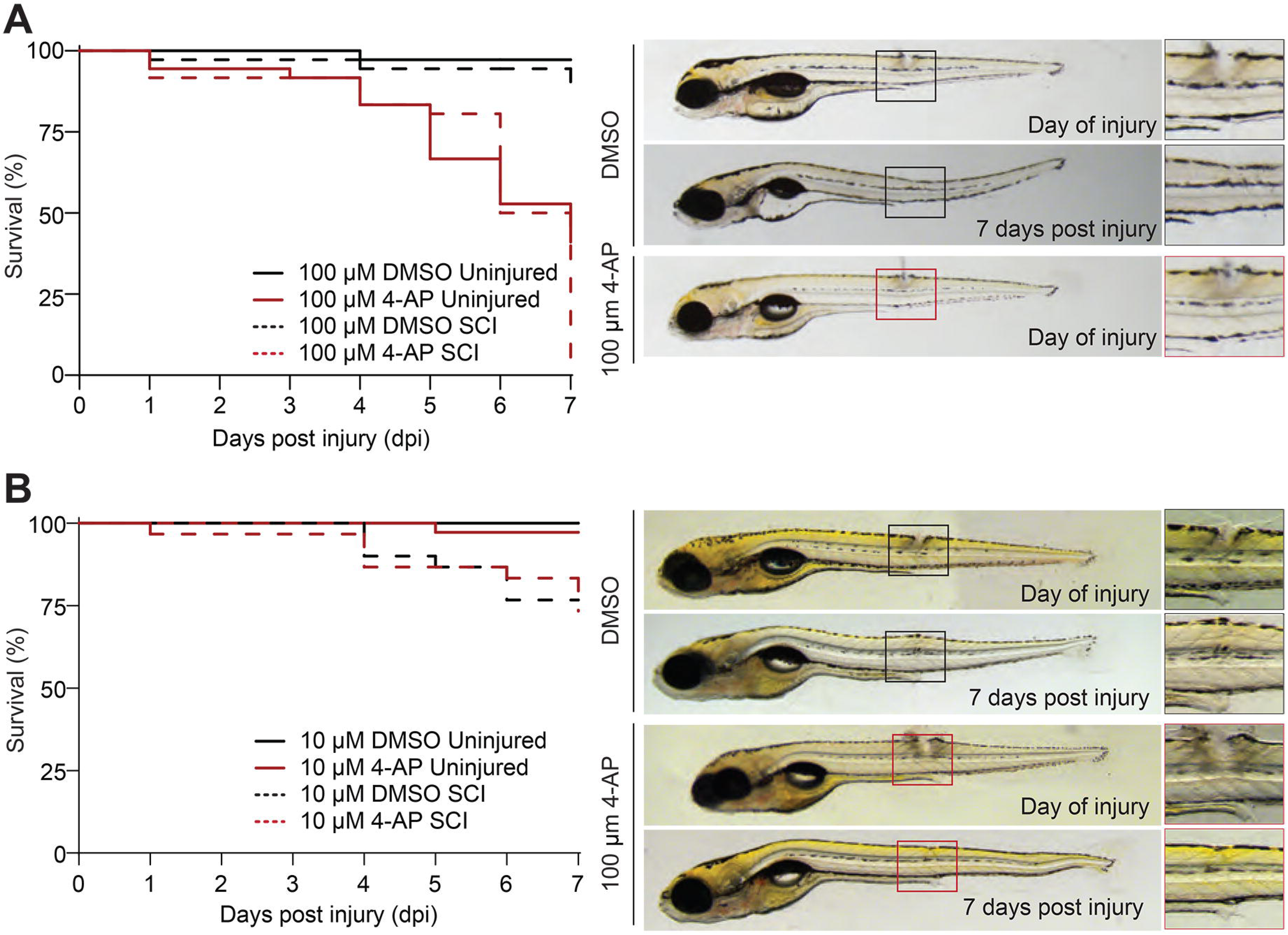
Toxicity of 4-AP in zebrafish larvae. Larvae were transected and survival measured for larvae treated with (**A**) 100 μM or (**B**) 10 μM 4-AP. Graphs are Kaplan-Meier plots for vehicle (black) and 4-AP (red) treated larvae. Larvae treated with either 100 μM or 10 μM 4-AP differed significantly from controls (χ^2^=15.37, p<0.0001 and χ^2^=22.85, p<0.0001). Representative micrographs of larvae are shown before and after one week exposure to vehicle or 4-AP. Insets highlight full recovery of injury in surviving larvae. Larvae orientation: lateral view, anterior left. (DMSO:n=60, 4-AP:n=62, DMSO+SCI:n=72, 4-AP+SCI:n=61)

**Figure 2.**
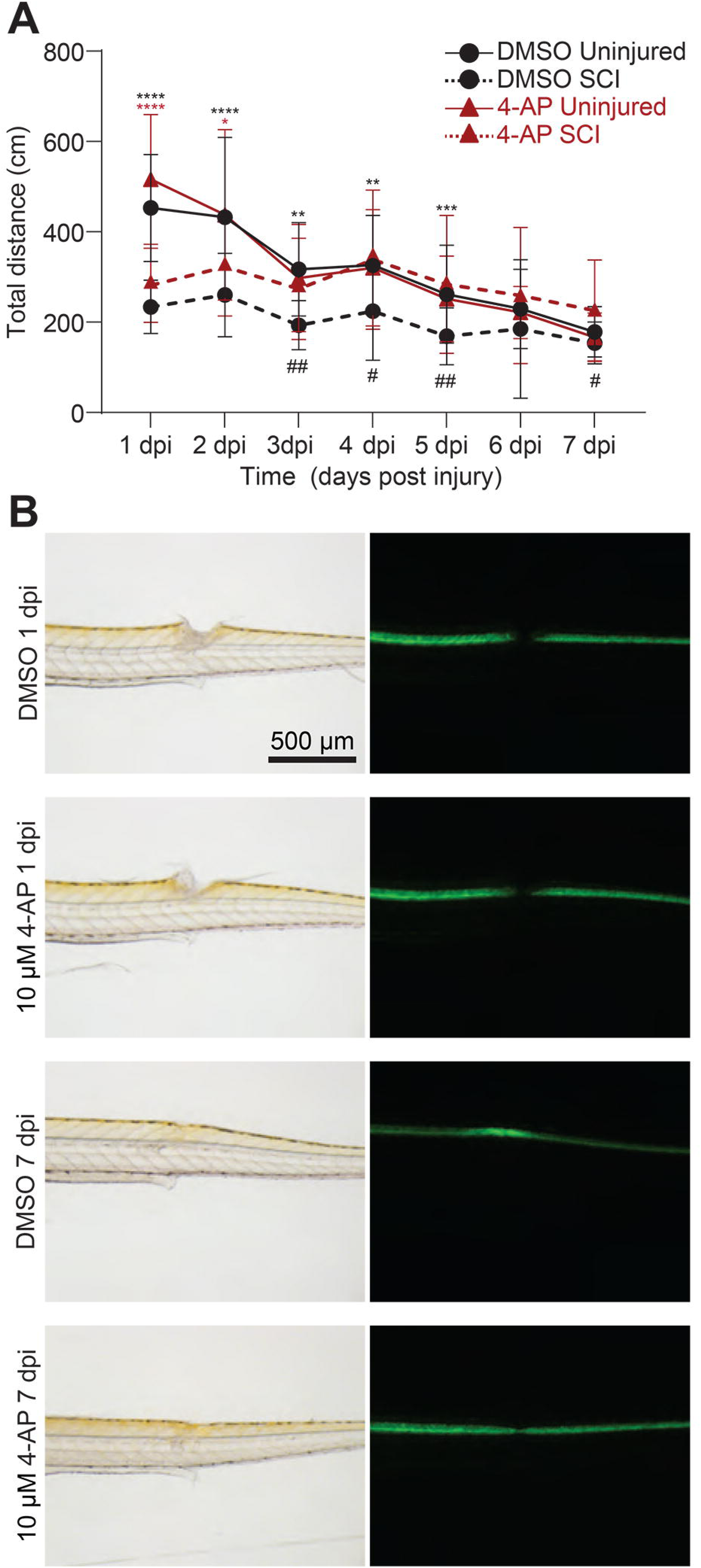
10 μM 4-AP enhances swim recovery and glial bridging after SCI. Individual larvae were treated with vehicle or 4-AP and tracked over 7 days. (**A**) Graph is average swim distance ± standard deviation. Asterisks denote significance between control and SCI within a treatment. Hashtags denote significance between SCI groups from a Tukey post-hoc test from a mixed-model ANOVA analysis with Geisser-Greenhouse correction for non-sphericity. (# p<0.05, **,## p<0.005, *** p<0.001, **** p<0.0001). (B) *Tg(gfap:EGFP)* larvae were transected at 5 days post fertilization and imaged at the beginning (1 dpi) and end (7 dpi) of experiment. Representative brightfield and fluorescence micrographs show full recovery of the tissue and formation of a glial bridge in green. Larvae orientation: lateral view, anterior left. (DMSO:n=36, 4-AP:n=35, DMSO+SCI:n=27, 4-AP+SCI: n=29).

**Figure 3.**
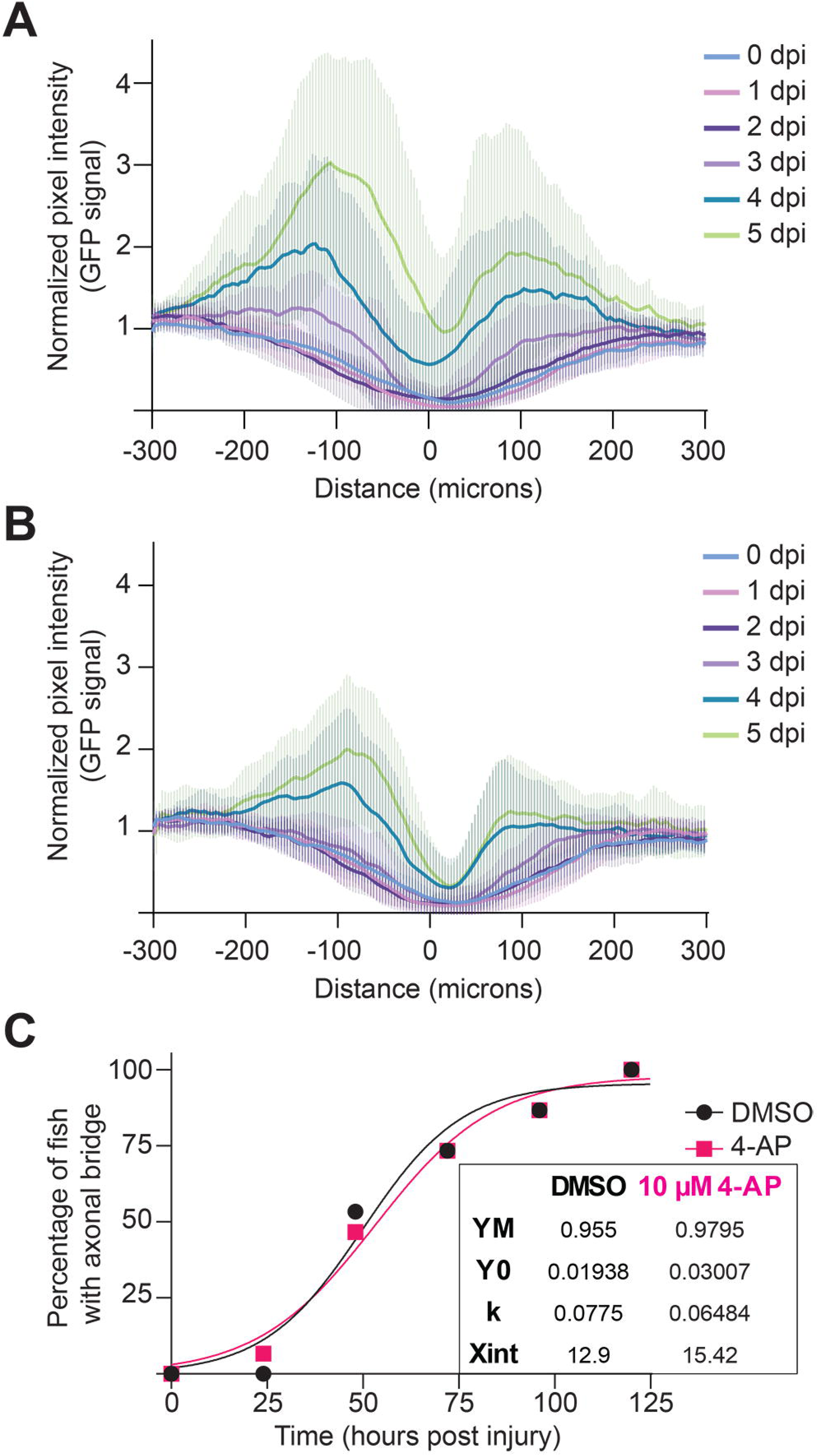
10 µM 4-AP decreases pixel intensity at the site of injury but does not affect the ability to form an axonal bridge. *Tg(gfap:EGFP; elavl3:mCherry-CAAX)* larvae were transected and followed individually for 5 days post injury (5 dpi). Quantification of mean pixel intensity (solid line) and standard deviation (shading) for transected larvae treated with (**A**) 0.1% DMSO and (**B**) 10 µM 4-AP graphed approximately every 10 μm for clarity. Epicenter of the lesion denoted as 0 on the X-axis was determined by the lowest average normalized pixel intensity value on day of injury (0 dpi). Pixel intensity increases proximal to the lesion and persists through bridge formation indicating a local increase GFP signal. (**C**) Percent of animals with an axonal bridge across the injury site was determined by complete tracks of fluorescence across the site of injury. Lines are the fit to a logistic growth curve with best fit values given. (DMSO+SCI:n=16, 4-AP+SCI:n=13)

**Figure 4.**
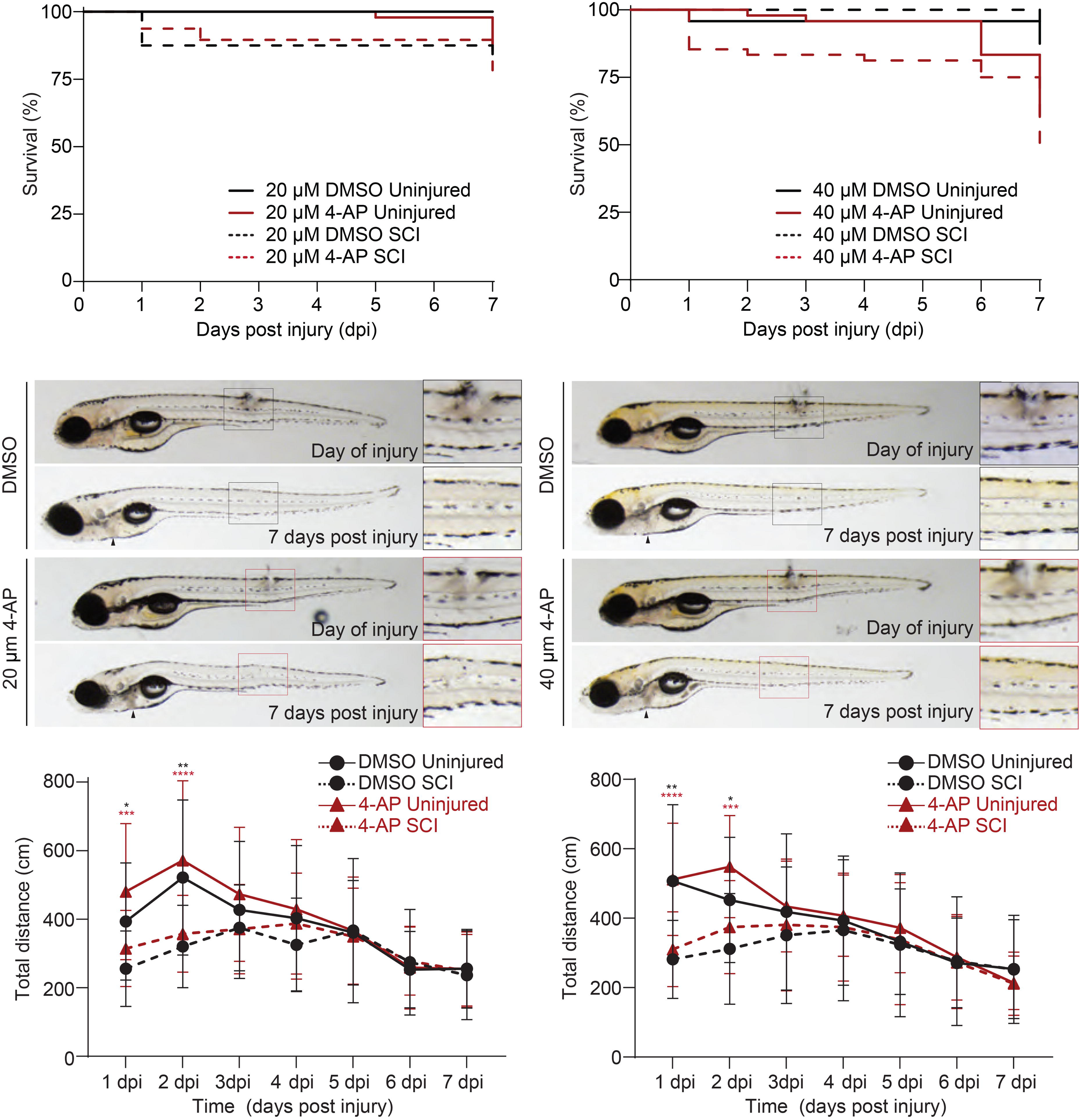
Higher concentrations of 4-AP do not enhance swim recovery. Larvae were either left uncut or transected and treated with either 20 μM or 40 μM 4-AP. (**A**) Survival graphs are Kaplan-Meier plots for vehicle (black) and 4-AP (red) treated larvae. (20 μM: χ^2^=6.500, p=0.0897 and 40 μM: χ^2^=18.65, p=0.0003) (DMSO:n=24, 4-AP:n=48, DMSO+SCI:n=24, 4-AP+SCI:n=48). (**B**) Representative micrographs of larvae are shown before and after one week exposure to vehicle or 4-AP. Insets highlight full recovery of injury in surviving larvae. Larvae orientation: lateral view, anterior left. Arrowheads denote the liver. (20 μM: DMSO:n=24, 4-AP:n=41, DMSO+SCI:n=20, 4-AP+SCI:n=37 40 μM: DMSO:n=21, 4-AP:n=29, DMSO+SCI:n=22, 4-AP+SCI:n=25) (**C**) Graph is average swim distance ± standard deviation. Asterisks denote significance between control and SCI within a treatment from a Tukey post-hoc test from a mixed-model ANOVA analysis with Geisser-Greenhouse correction for non-sphericity. (* p<0.05, ** p<0.005, *** p<0.001, **** p<0.0001). (20 μM: DMSO:n=24, 4-AP:n=41, DMSO+SCI:n=20, 4-AP+SCI:n=37 40 μM: DMSO:n=21, 4-AP:n=29, DMSO+SCI:n=22, 4-AP+SCI:n=25).

