## Supplementary figures and images for "4-Aminopyridine Produces Concentration-Dependent Improvement in Locomotor Recovery after Spinal Cord Injury in Zebrafish Larvae^1^"

### Supp Figure 1

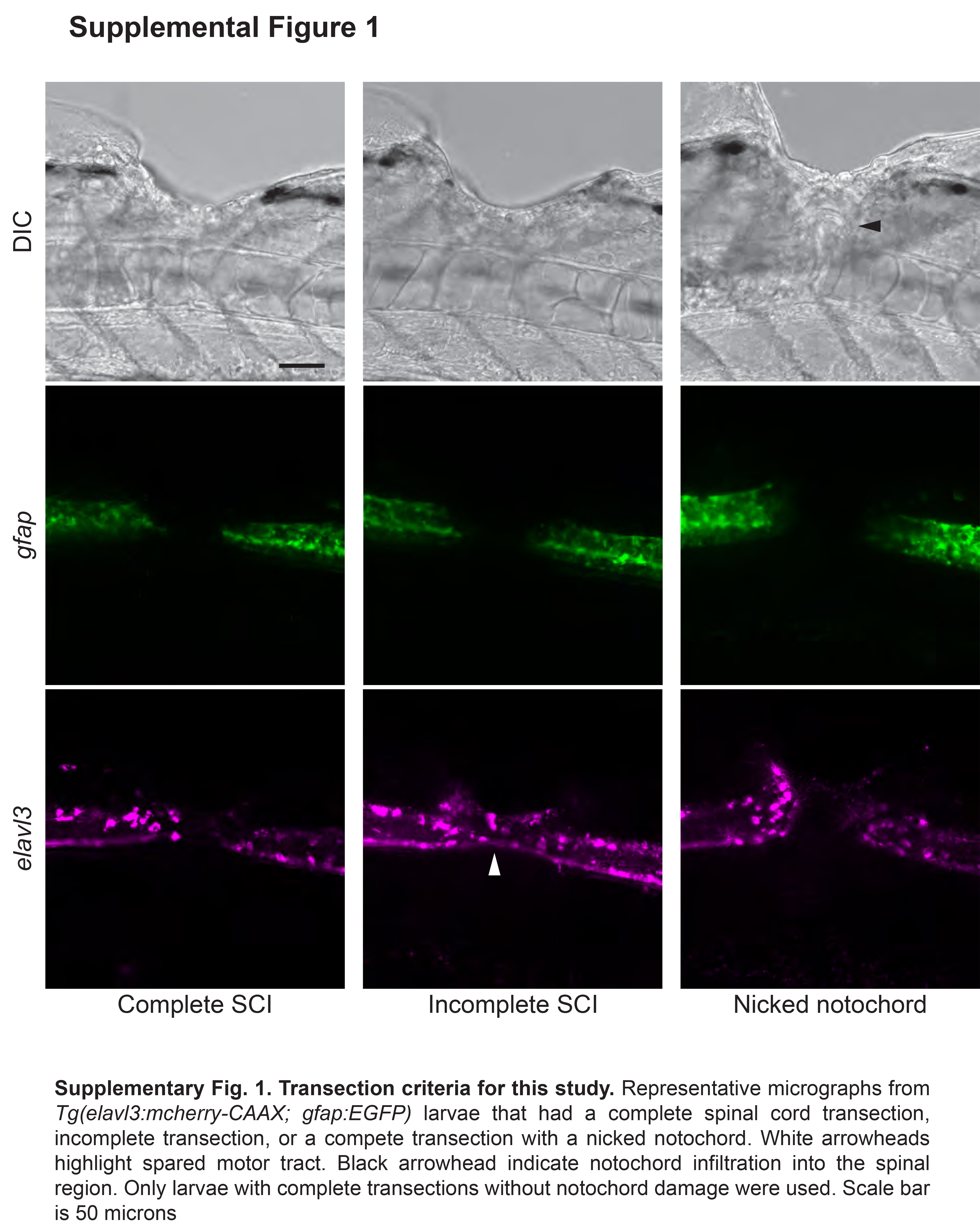

### Supp Figure 2

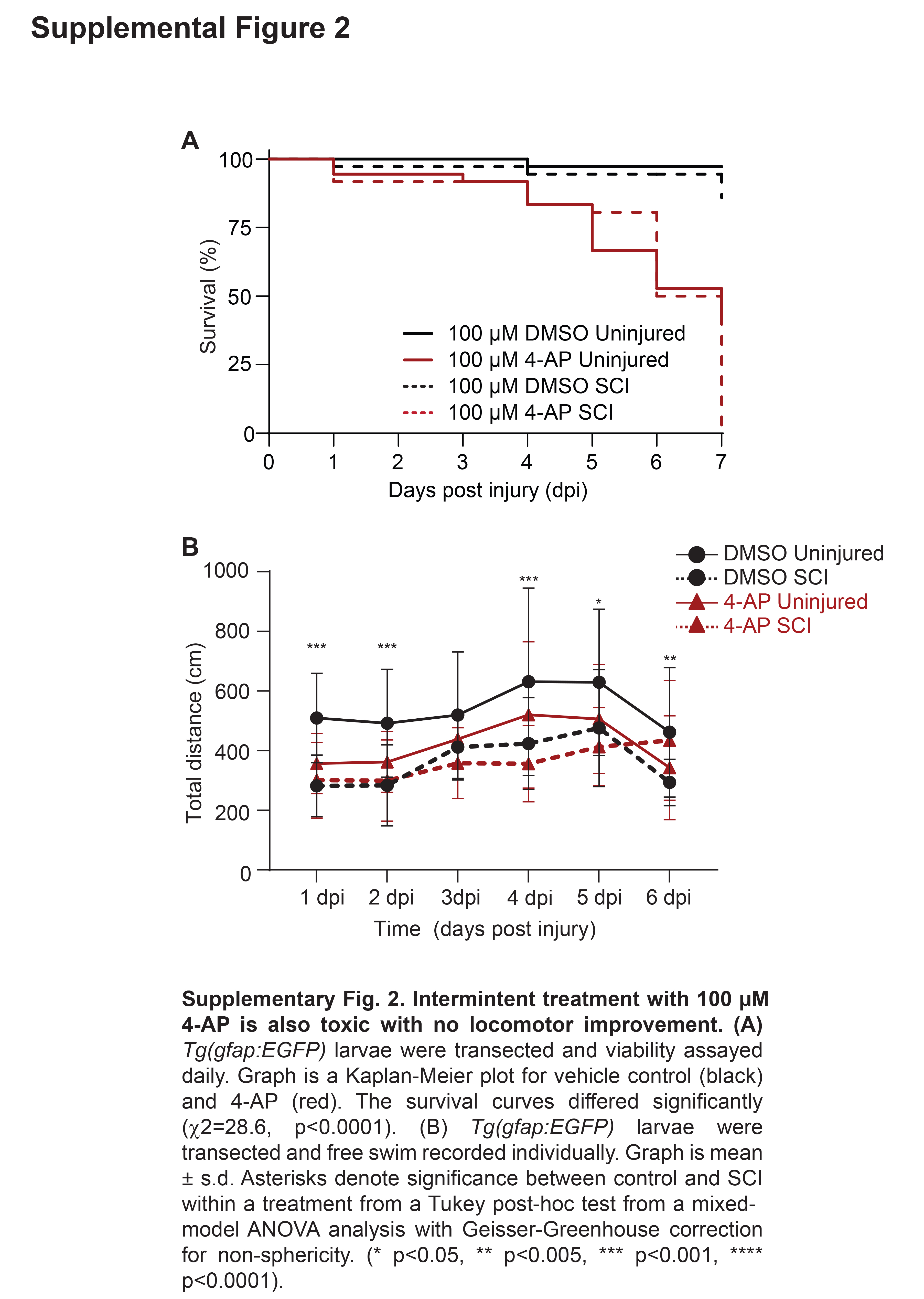

### Supp Figure 3

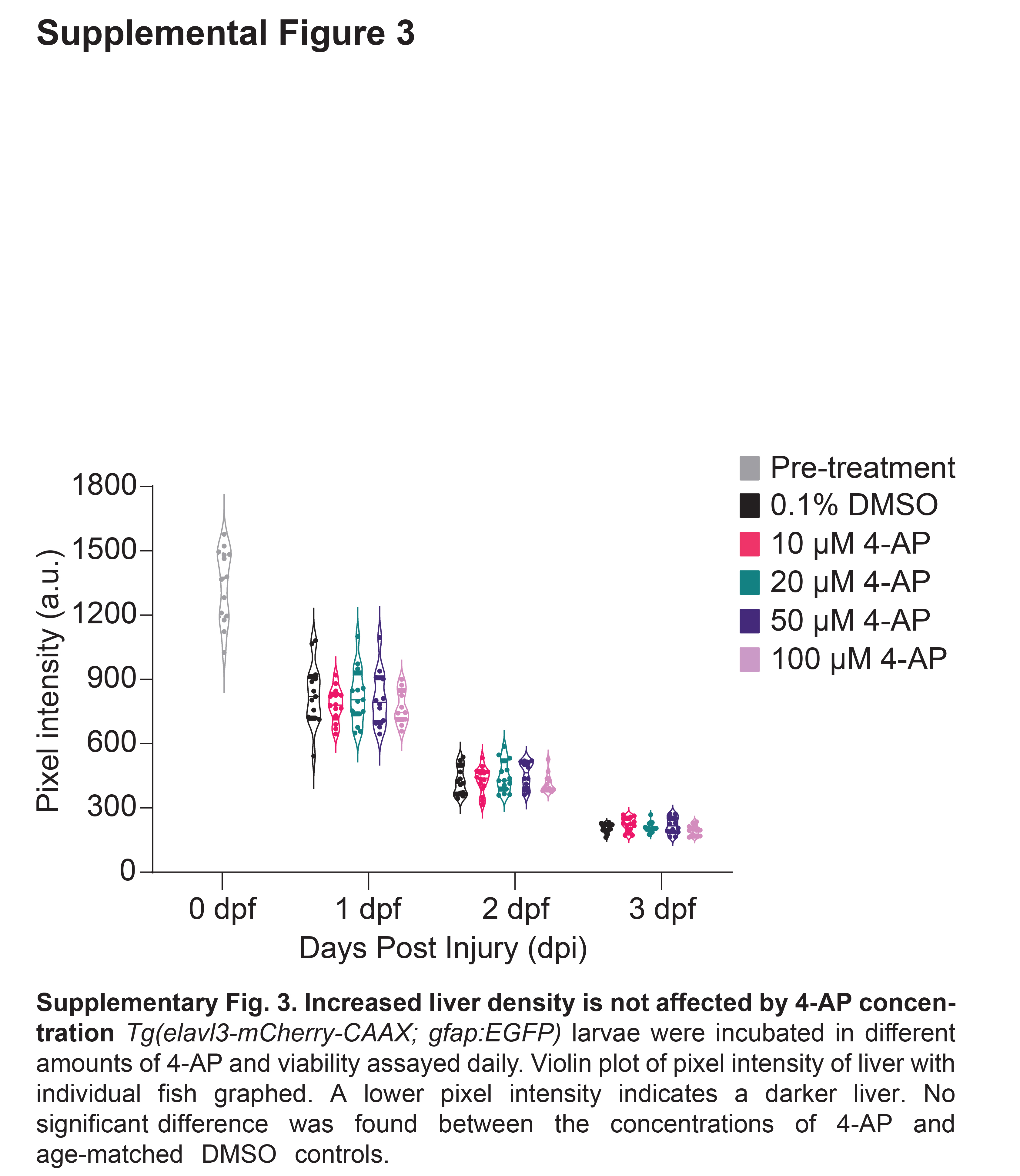
